# High Refractive Index Imaging Buffer for Dual Color 3D SMLM Imaging of Thick Samples

**DOI:** 10.1101/2024.04.30.591648

**Authors:** Lulu Zhou, Wei Shi, Shuang Fu, Mengfan Li, Jianwei Chen, Ke Fang, Yiming Li

## Abstract

The current limitations of single-molecule localization microscopy (SMLM) in deep tissue imaging, primarily due to depth-dependent aberrations caused by refractive index (RI) mismatch, present a significant challenge in achieving high-resolution images at greater depths. To extend the imaging depth, we optimized the imaging buffer of SMLM with RI matched to the objective immersion medium, and systematically evaluated five different RI matched buffers, focusing on their impact on the blinking behavior of red-absorbing dyes and the quality of reconstructed super-resolution images. Particularly, we found clear unobstructed brain imaging cocktails (CUBIC) based imaging buffer could match the RI of oil and was able to clear the tissue samples. With the help of RI matched imaging buffer, we showed high quality dual color 3D SMLM images with imaging depth ranging from few microns to tens of microns in both cultured cells and sectioned tissue samples. This advancement offers a practical and accessible method for high-resolution imaging at greater depths without specialized optical equipment or expertise.

## Introduction

Single-molecule localization microscopy (SMLM) has been widely used to dissect how target molecules are arranged in three dimensions (3D) with unprecedented resolution (*1-3*). However, this powerful technique encounters a critical optical limitation when applied to thicker samples. The primary obstacle is the refractive index (RI) mismatch between the sample and the objective lens’ immersion medium, a discrepancy that results in depth-dependent aberrations, thus blurring and distorting images at increased depths (*4, 5*). For super-resolution imaging, biological samples are typically embedded in water-based buffer and imaged with a high numerical aperture (NA) oil immersion objective (*6*). Although high NA objectives could collect more photons and form a sharper point spread function (PSF), the depth-dependent aberrations restrict the imaging depth of SMLM to few microns close to the coverslip.

To extend the imaging depth (i.e., tens of microns) for SMLM, adaptive optics (AO) was used to compensate for the aberrations (*7-9*). However, the requirement of dedicated optics (e.g., deformable mirror) and expertise limited the implementation of AO only in few expert labs. An alternative method is to match the RI between the immersion and mounting medium. RI matched buffer has been widely used in thick sample imaging such as multiphoton microscopy, light sheet microscopy, and optical clearing (*10-14*). By matching the refractive indices, not only aberrations could be reduced, but also the scattering of the light would be suppressed. In contrast to the conventional RI matched buffer where only traditional optical properties (i.e., RI, absorption/emission spectra) are evaluated, the blinking properties of the fluorophores need to be additionally evaluated for RI matched buffer used for SMLM imaging. Huang et al., first introduced glycerol and sucrose-based buffers for SMLM imaging which could reach refractive index of 1.45 to alleviate the depth-dependent aberration (*15*). We also introduced 2,2’-thiodiethanol (TDE), a nontoxic hydro soluble reagent with high RI (1.5215) (*16*), to the primary thiol and oxygen scavenger based imaging buffer to match the RI (1.406) of silicone oil (*17*). Recently, Kim lab reported a SMLM imaging buffer with RI matched to the oil (RI = 1.518) by adding the hydro soluble reagent, 3-pyridinemethanol (3-PM, RI = 1.545) (*18*). However, all the imaging buffers were only demonstrated in cultured thin single layer cells. RI matched buffer for super resolution imaging of thicker samples such as tissue sections is still missing in the field.

Here, based on the whole-brain clearing method CUBIC (clear unobstructed brain imaging cocktails) (*19*), we introduced a hydrophilic chemical mixture (antipyrine and nicotinamide) to SMLM imaging buffer, called CUBIC-R+, to match the RI of oil. We evaluated various RI matched buffers (i.e., TDE, glycerol, sucrose, 3-PM and CUBIC-R+ based imaging buffer), tailored for different types of objectives (i.e., oil and silicone oil objectives), and rigorously tested their compatibility with SMLM. We systematically assessed the buffers’ impact on the performance of two commonly used red blinking dyes, Alexa Fluor 647 (AF647) and CF680. Photostability and blinking behavior of these dyes within the RI matched buffers were quantified. We verified the compatibility of 3D and dual color SMLM imaging with the RI matched buffers by benchmark of the referenced structure proteins (nuclear pore complex protein Nup96 for 3D, microtubule protein β-Tubulin and mitochondria outer membrane protein Tom20 for dual color). Furthermore, we performed 3D SMLM imaging of the biological samples through mouse brain slices using CUBIC-R+ buffer and reconstructed high quality 3D super resolution images without obvious distortion, demonstrating its ability for both tissues clearing and SMLM imaging.

## Results

### Depth-dependent PSFs

We first recorded the z-stack images of beads with oil objective for observing the 3D astigmatic PSFs. To directly reveal the effect of RI mismatch between mounting medium and the oil (RI = 1.518) on PSF, two mounting media, MilliQ water (RI = 1.33) and CUBIC-R+ (RI = 1.518) were used to record the 3D PSFs at different depths (0-30 μm), respectively. As shown in Fig. 1a, samples of beads were mounted between two pieces of cover glass as a sandwich with the thickness adjusted by a spacer. Comparing the PSFs at the bottom and upper surface, we found the PSF in water varied with depth whereas the PSF in CUBIC-R+ did not (Fig. S1). We evaluated the lateral and axial resolutions at different depths by measuring the corresponding full width at half maximum (FWHM) of PSF. As shown in Fig. 1b and 1c, the resolutions estimated by PSF from CUBIC-R+ did not change with the increasing depth while the resolutions, especially for the axial resolution, estimated by PSF from water deteriorated with the increase of depth. Similar results were observed for 3D PSFs at different depths in MilliQ water and TDE based RI matched buffer imaged with silicone oil (RI = 1.406) objective (Fig. S2). Altogether, we verified the mismatch between RIs in mounting medium and objective immersion medium could cause the depth-dependent aberration, which deteriorated the resolutions, especially for the axial resolution at the deeper depth. Another benefit for RI matching is that the experimental PSF measured on the cover glass surface can be directly applied to analyze the data deep inside the sample with minimum distortions.

**Fig. 1.**
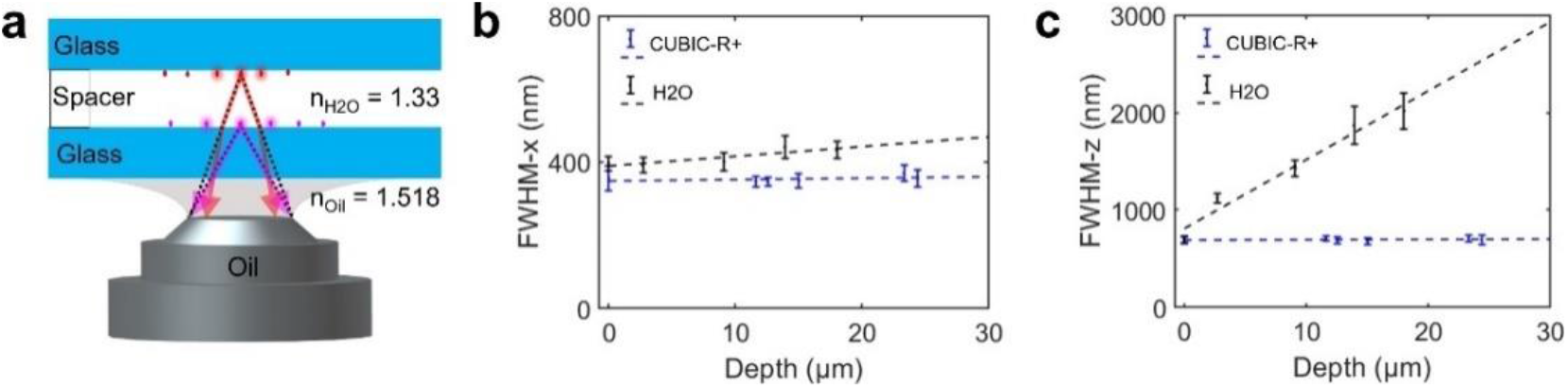
Quantification of the resolution at different focusing depth with beads. (a) Diagram of experimental setup for evaluating depth-dependent 3D PSFs. The thickness between beads on two cover glasses were adjusted by the volume of mounting medium (H2O or CUBIC-R+) or the addition of brain slice (20 μm thick) as a spacer. (b, c) Lateral (b) and axial (c) FWHM of PSF as a function of the medium and depth. The lateral and axial FWHM in RI matched medium (CUBIC-R+) remain constant with increasing depth, whereas axial FWHM in H2O increased with the increase of depth. The lateral FWHM in H2O just increased slightly with the increase of depth.

### Photoswitching properties of AF647 and CF680 in RI matched imaging buffer

We mixed the high RI chemical reagents with the conventional water-based SMLM imaging buffer, called as water-index (WI) buffer for short, to reach the desired RI (oil 1. 518 or silicone oil 1.406). Considering that the imaging buffer composition played a vital role in the photoswitching properties of dyes for SMLM (*20, 21*), we then selected two red-absorbing dyes, AF647 and CF680 (most commonly used dye pairs for ratiometric dual-color imaging) (*22*), as benchmarks, and determined the impact of the substantial RI matched reagents on the blinking behavior of the fluorophores. Here, two oil-index buffers (CUBIC-R+ and 3-PM) and three silicone oil-index buffers (TDE, glycerol and sucrose) were tested while the WI imaging buffer was selected as control.

To evaluate the photoswitching characteristics, four parameters including number of photons per switching event, number of switching cycles, on-off duty cycle (DC) and survival fraction (SF) was determined from the single-molecule fluorescence time traces (*23, 24*). As shown in Table 1, we summarized the switching properties of dyes in RI matched imaging buffers. For oil-index buffers, we found both AF647 and CF680 in CUBIC-R+ had higher photons and photostability than those in 3-PM (Fig. S3 and S4). Though the photons in oil-index buffers decrease in contrast with that in WI buffer, both AF647 and CF680 dyes exhibit high enough photons and equal low DC values in oil-index buffers, suitable for SMLM imaging. Besides, we found that 3-PM showed light yellow color while CUBIC-R+ was colorless and less viscous than 3-PM (Fig. S5). Due to the different appearance between CUBIC-R+ and 3-PM, we tested the absorption and fluorescence spectra and compared the optical properties for the two buffers. As shown in Fig. S6, different from the 3-PM buffer which has a strong absorption below the wavelength of 450 nm, CUBIC-R+ has much less absorption. In line with the absorption, we also found CUBIC-R+ has a much weaker emission than 3-PM (Fig. S6) when excited with laser of different wavelengths (405, 488, 561 and 642 nm). These results suggested that CUBIC-R+ oil-index buffer had superior optical properties and blinking behaviors than 3-PM. Comparing the silicone oil-index buffers with WI buffer, both AF647 and CF680 dyes show the comparable high (or slightly improved) photons and adequate low DC values (Fig. S7 and S8), which demonstrates the silicone oil-index TDE/glycerol/sucrose-based buffers are also well compatible with SMLM imaging.

**Table 1.** Photoswitching properties of AF647 and CF680 in different buffers imaged by an oil or silicone oil objective.

| Dye | objective | Buffer | Photons per switching event | Number of switching cycles (mean) | Equilibrium on-off duty cycle (400-600 s) | Survival fraction after illumination for 400 s |
| --- | --- | --- | --- | --- | --- | --- |
| AF647 | Oil<br>NA = 1.5 | CUBIC-R+ | 6334±854 | 10±1 | 0.0005±0.0001 | 0.79±0.02 |
|  |  | 3-PM | 4570±722 | 15±1 | 0.0005±0.0001 | 0.68±0.02 |
|  |  | WI | 13693±2312 | 13±2 | 0.0004±0.0001 | 0.81±0.16 |
|  | Silicone oil<br>NA = 1.35 | TDE | 8282±448 | 9±1 | 0.0004±0.0001 | 0.81±0.04 |
|  |  | Gly | 10387±1442 | 7±1 | 0.0004±0.0001 | 0.77±0.03 |
|  |  | Suc | 10306±1991 | 9±2 | 0.0004±0.0001 | 0.78±0.05 |
|  |  | WI | 8283±1812 | 10±2 | 0.0004±0.0001 | 0.80±0.05 |
| CF680 | Oil<br>NA = 1.5 | CUBIC-R+ | 5669±506 | 27±7 | 0.0006±0.0002 | 0.82±0.04 |
|  |  | 3-PM | 4283±356 | 15±1 | 0.0005±0.0001 | 0.68±0.03 |
|  |  | WI | 7559±582 | 23±3 | 0.0007±0.0001 | 0.79±0.07 |
|  | Silicone oil<br>NA = 1.35 | TDE | 4005±527 | 30±4 | 0.0004±0.0002 | 0.76±0.06 |
|  |  | Gly | 4344±704 | 33±5 | 0.0007±0.0001 | 0.80±0.03 |
|  |  | Suc | 4425±785 | 39±7 | 0.0011±0.0006 | 0.71±0.04 |
|  |  | WI | 3688±221 | 34±8 | 0.0010±0.0005 | 0.82±0.06 |

### Cellular imaging of nuclear pore complex

We then evaluated the performance of the RI matched buffer by 3D SMLM imaging of Nup96 which is a scaffold nucleoporin of human nuclear pore complex (NPC) and often served as a reference protein due to its stereotypical geometry (*25, 26*). For WI buffer, the double ring structure of the NPC can be nicely resolved in the bottom surface of nuclear envelopes (NEs) which is close to the coverslip (Fig. 2d and 2e). However, the double ring structure can hardly be resolved in the upper surface of NEs due to the spherical aberration caused by the RI mismatch (Fig. 2c and 2e). Even for the lower surface of the NEs, we found that there is a strong correlation between the double ring distance and the axial position of the NPCs (Fig. 2f). This suggested that the refractive index mismatch could distort the super resolution image even in the range of a few hundred nanometers close to the coverslip. We then used the oil-index buffers to image Nup96. Distinct from WI buffer, both CUBIC-R+ and 3-PM could nicely resolve the two rings with eight corners on both lower and upper surface of NEs (Fig. 2a, 2b, 2e and S9), indicating that the spherical aberration was significantly reduced. There was also no correlation between the double ring distance and the axial position of the NPCs (Fig. 2f and S9d), indicating that depth-dependent aberrations were minimized. To further quantify the image quality in oil-index buffers, we determined the photon number, effective labeling efficiency (ELE) and Fourier ring correlation (FRC) resolution for super-resolution images of Nup96. We found comparable photons for CUBIC-R+ and WI buffers measured in cellular environment, while the photons for CUBIC-R+ is much less than WI measured on cover glass (Table 1 and Fig. 2g). It highlights that high RI buffer could collect more photons when a high NA objective is used. The ELE for CUBIC-R+ is also adequate high as WI (Fig. 2g and S9f). The FRC resolution is close for CUBIC-R+ (16.4 nm) and WI (19.8 nm) buffers-based imaging, while it decreases to 28.1 nm for 3-PM buffer-based imaging (Fig. 2h). By plotting the number of localizations per volume as a function of time, we found the fluorescent signal of Nup96 in 3-PM decayed far more rapidly than that in CUBIC-R+ (Fig. S9g), which agreed with the single dye molecule experiment on the coverslip where the survival fraction of the dyes in CUBIC-R+ is higher than that in 3PM (Table 1). These results suggested that CUBIC-R+ exhibited a superior performance in SMLM imaging than 3-PM.

**Fig. 2.**
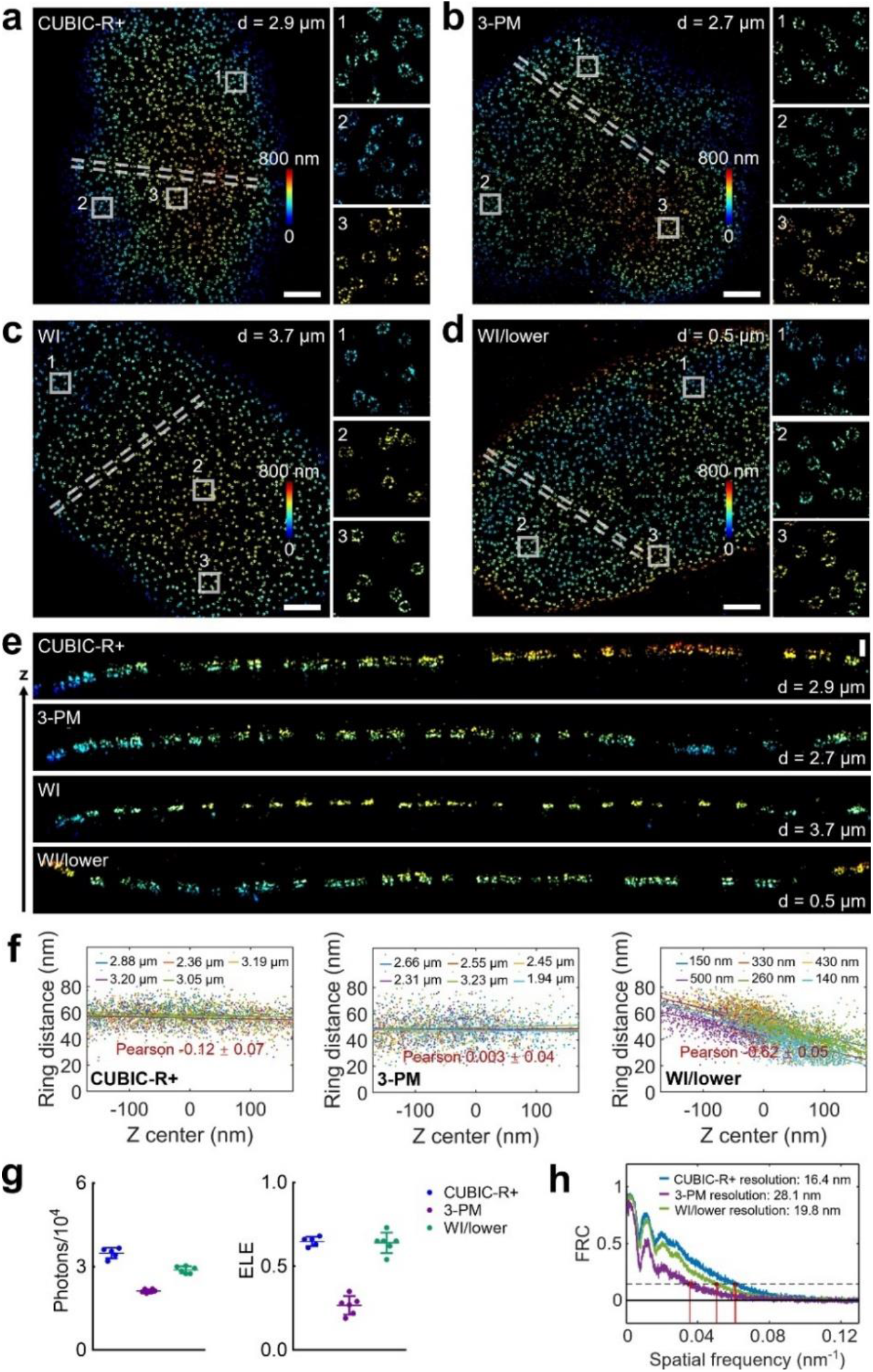
Evaluation of the influence of oil-index buffers on 3D SMLM imaging of NPCs. (a-c) Representative images of Nup96 proteins on the upper nuclear envelope in CUBIC-R+ (a), 3-PM (b) and WI (c) buffers. The right panels show zoomed images for regions boxed by white square, and the color bars represent the z position. (d) Representative image of Nup96 on the lower nuclear envelope in WI buffer. (e) The cross-sectional view images of the white dashed regions in (a-d). (f) Plots of distance between two rings versus the z position for Nup96 in different depths and buffers. (g) Scatter plots of the photon number (left) and ELE (right) for Nup96 in different buffers. Each symbol corresponds to one cell nucleus. Horizontal bars and errors show the means and SD, respectively. (h) FRC curves of Nup96 in regions corresponding to the left panels of (a, b and d). Scale bars: 2 μm (a-d), 200 nm (e).

Analogously, we tested the performance of silicone oil-index buffers in the 3D SMLM images of Nup96 on lower or upper NEs with a silicone oil objective (Fig. S10 and S11). Using silicone oil-index buffers, the two rings for Nup96 on both lower and upper NEs could be nicely resolved (Fig. S10e and S11e). There is also no correlation between the ring distance and z positions of NPCs (Fig. S10f and S11f). In comparison with WI buffer-based imaging of Nup96, we found the ELE values for silicone oil-index buffers-based imaging are comparable (Fig. S10g and S11g). Comparing with the FRC resolution for WI buffer-based imaging, we also found the comparable high (or slightly improved) resolutions for silicone oil-index buffers-based imaging (Fig. S10h and S11h), which demonstrates the silicone oil-index TDE/glycerol/sucrose-based buffers are also well compatible with SMLM imaging of the substructures in a cell.

### Dual-color imaging of microtubules and mitochondria

We then evaluated the performance of the RI matched buffer in the dual-color 3D SMLM imaging experiments. Immunostained microtubules (CF680) and mitochondria (AF647) were imaged with ratiometric 3D SMLM. Close to the coverslip, we found all the oil-index or silicone oil-index buffers had similar reconstructed dual color 3D super-resolution images as that of WI buffer, nicely separating the filamentous microtubules (MTs) from the globular or tubular mitochondria (Fig. 3a-d and S12). The corresponding side views (Fig. 3d and S12e), near the coverslip, also clearly revealed the elliptical hollow structure of outer mitochondrial membranes (OMMs) with the width of minor axis ranging from 250 to 500 nm. The separation of the adjacent MTs in z and the contacts between MTs and OMMs were also well resolved. Analyzing the FRC resolution for microtubules and mitochondria (Fig. 3e and 3f), we found comparable resolution for CUBIC-R+ (21.0/29.1 nm) and WI (23.7/29.7 nm) based buffers, whereas better than that for 3-PM (27.9/35.1 nm) based buffer.

**Fig. 3.**
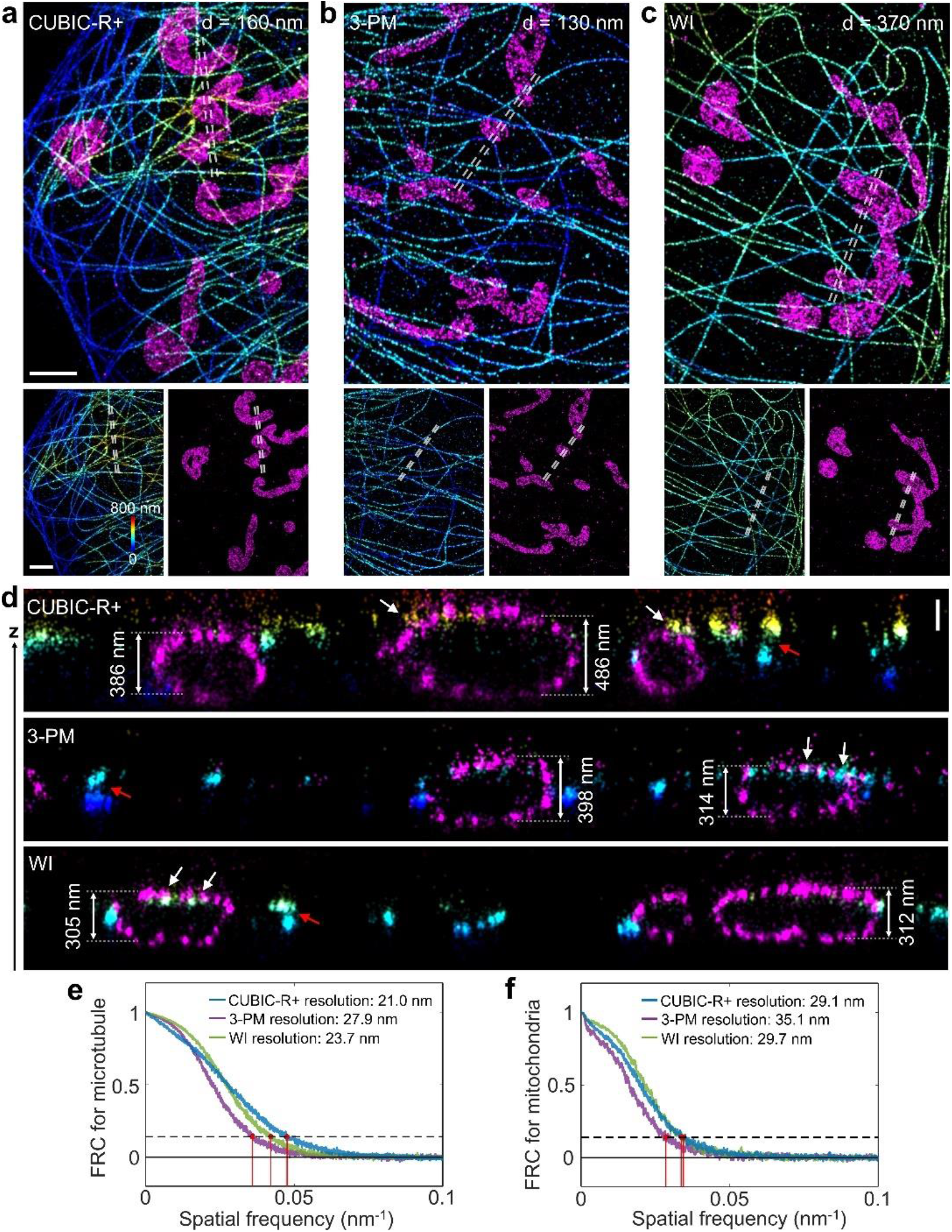
Dual-color 3D SMLM imaging of microtubules and mitochondria slightly above the basal membrane of COS-7 cells in oil-index or WI buffers. (a-c) Representative images of microtubules (anti-β-tubulin/CF680) and the outer mitochondrial membrane (anti- Tom20/AF647) in CUBIC-R+ (a), 3-PM (b) and WI (c) buffers. Color bar denotes the z position. (d) The side view of the white dashed regions in (a-c). The white arrows indicate the mitochondria-microtubule contacts, while the red arrows denote the different layers of microtubule filaments. (e, f) FRC curves of microtubule (e) or mitochondria (f) in regions corresponding to the bottom panels of (a-c). Scale bars: 2 μm (a-c), 200 nm (d).

We then performed dual-color 3D SMLM deep in the cell (∼ 10 μm) using oil objective. CUBIC-R+ and WI based buffers were used as RI matched and control buffers, respectively. In CUBIC-R+, the representative dual-color images at the depth of 11.1 μm (Fig. 4a) clearly showed the MT filaments and the mitochondrial morphology as a tubular and interconnected network. The separation for axial adjacent MTs, the mitochondria-microtubule contacts and the elliptical hollow structure for OMMs were also nicely resolved in the cross-sectional view, which is similar to those near the coverslip (Fig. 4b). However, in WI based buffer, the dual-color 3D images of MTs and OMMs at the depth of 6.4 μm (Fig. 4c and 4d) showed squeezing and distortions in z. The width of the mitochondria along z axis in WI based buffer was compressed. The MT filaments appeared flat in z, without obvious z variation between different filaments. Moreover, it was difficult to accurately resolve the mitochondria-microtubule contacts from the dual-color imaging in WI based buffer (Fig. 4d). These results further suggested the necessary of applying RI matched buffers in deep 3D SMLM imaging, since RI matched buffers could evade the PSF model mismatch and localization errors induced by depth-dependent aberrations.

**Fig. 4.**
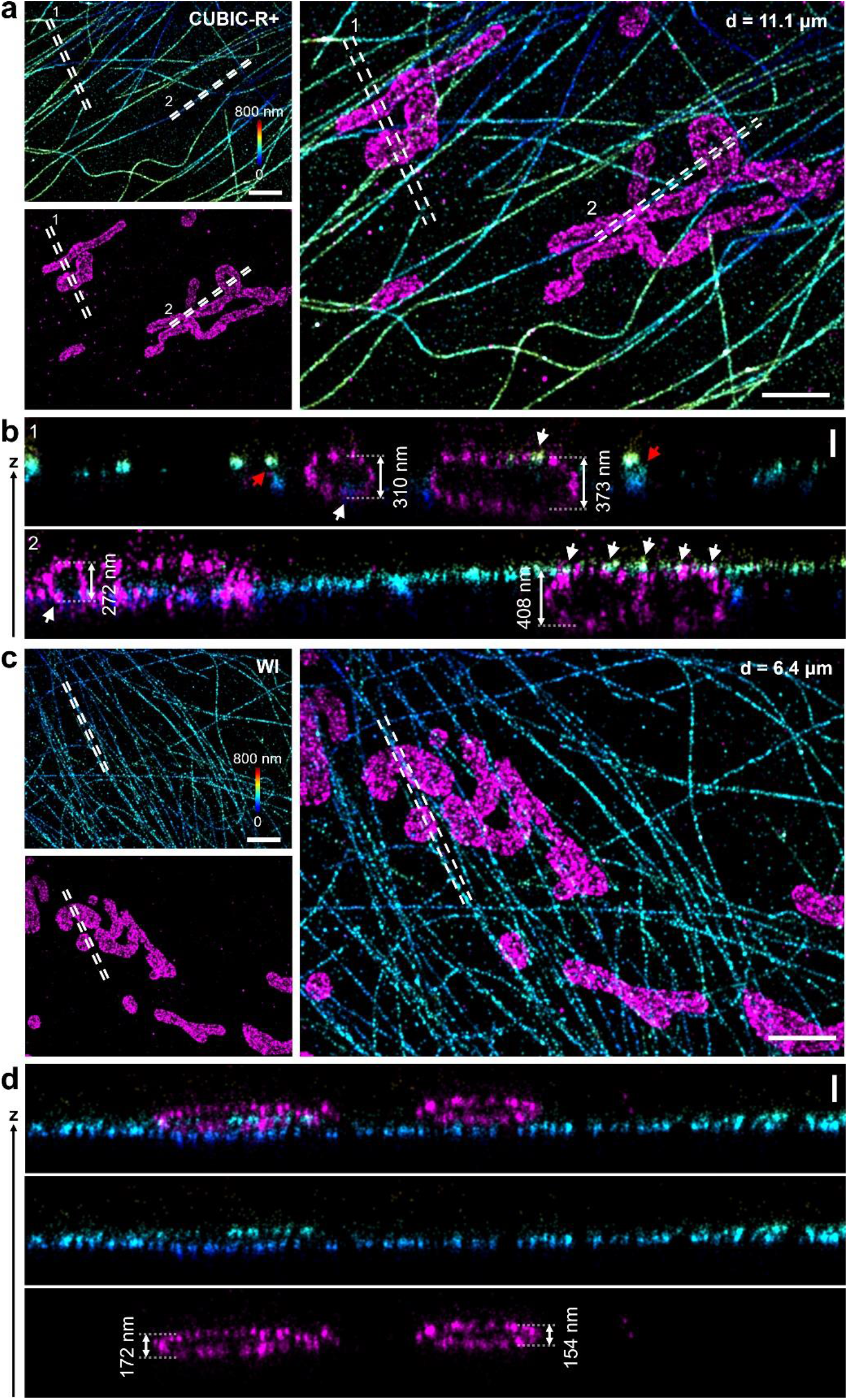
Comparison of the deep imaging of microtubules and mitochondria in CUBIC-R+ and WI buffers by placing COS-7 cells upside down on a coverslip. (a, c) Dual-color 3D SMLM images of microtubules (anti-β-tubulin/CF680) and the outer mitochondrial membrane (anti-Tom20/AF647) in CUBIC-R+ (a) and WI (c) buffers. Color bars denote the z position. (b, d) The side view of the white dashed regions in (a, c). The white arrows indicate the mitochondria-microtubule contacts, while the red arrows denote the different layers of microtubule filaments. Scale bars: 2 μm (a, c), 200 nm (b, d).

### Deep imaging of cellular structure on tissue

Finally, we applied the RI matched buffer to the more challenging tissue sample which was several tens of microns away from the coverslip and imaged with an oil objective. Since CUBIC-R+ was used for tissue clearing previously (*19, 27*), we conducted the clearing experiments by immersing the mouse brain tissue slices (∼ 30 μm thick) in CUBIC-R+ buffer for 10 min before taking photos. As shown in Fig. 5a, the sagittal brain tissue in CUBIC-R+ based imaging buffer was almost transparent, whereas the tissue in 3-PM based buffer remained some opaque regions. To test the capability of CUBIC-R+ in tissue imaging, we first imaged the reference Nup96 proteins by placing the U2OS-Nup96-SNAP cells on top of a mouse brain tissue (Fig. 5b). With CUBIC-R+, the double ring structure of NPC can still be resolved at the depth of ∼26.4 μm (Fig. 5c and 5e), whereas the double ring structures for Nup96 could not be clearly distinguished at similar depth with 3-PM (incomplete rings, Fig. 5d and 5e). There was also no correlation between the ring distance and z positions of the NPCs, indicating depth-dependent aberrations can be corrected even for thick tissue samples with our RI matched buffer (Fig. 5f). The photon number and ELE values were also similar as those in the samples near coverslip (Fig. 2g and Fig. 5g).

**Fig. 5.**
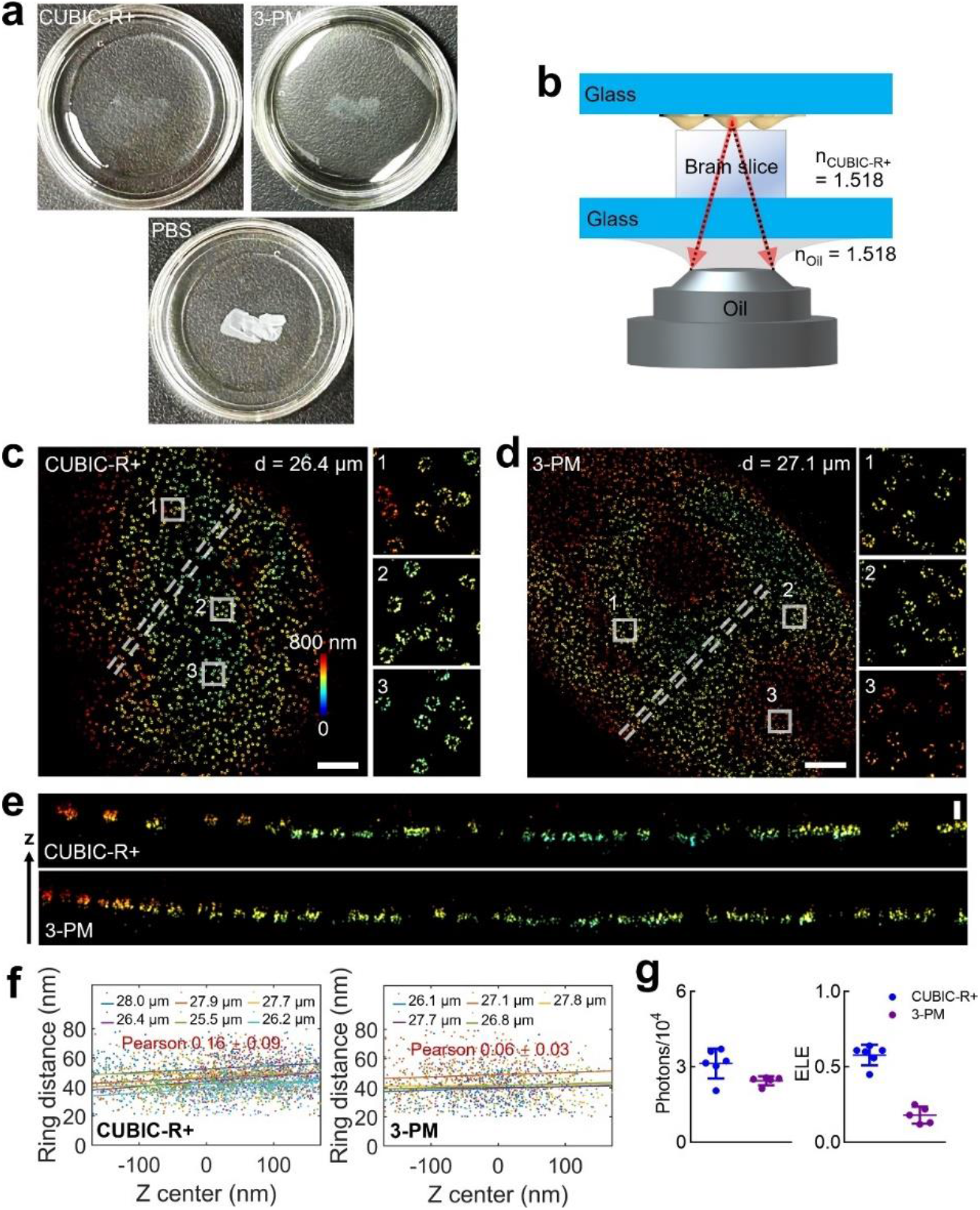
Deep imaging of NPCs in oil-index buffers by placing U2OS-Nup96-SNAP cells upside down on a fixed brain tissue slice. (a) Three mouse brain tissue slices (30 μm thick) were imbedded in CUBIC-R+, 3-PM and PBS for 10 min before taking photos, respectively. (b) Diagram of experimental setup for deep imaging of NPCs. (c, d) 3D SMLM images of Nup96 labeled by AF647 in CUBIC-R+ (c) and 3-PM (d) buffers. The right panels show zoomed images for regions in white, and the color bar show the z position. (e) The side view images of the white dashed regions in (c, d). (f) Plots of distance between the rings versus the z position for Nup96 in different depths and buffers. (g) Scatter plots of the photon number (left) and ELE (right) for Nup96 in oil-index buffers. Each symbol corresponds to one cell nucleus. Horizontal bars and errors show the means and SD, respectively. Scale bars: 2 μm (c, d), 200 nm (e).

We further tested the compatibility of CUBIC-R+ buffer with the thick tissue samples by imaging cellular structure inside the mouse brain tissue. Here, myelin oligodendrocyte glycoprotein (MOG), a central nervous system myelin-specific protein, which preferentially localizes at the outermost surface of myelin sheaths or the extracellular surface of oligodendrocytes (Ols) (*28, 29*), was imaged. By immunofluorescent staining, we recorded the 3D SMLM images of MOG proteins in CUBIC-R+ at depth of ∼ 20 μm (Fig. S13), which clearly showed the local spatial distribution of Ols in the coronal brain section. Considering the compatibility with SMLM imaging and the ability to clear tissue samples, CUBIC-R+ buffer is an ideal candidate for tissue imaging using high NA oil objective.

## Discussion

In summary, we comprehensively optimized and characterized imaging buffers for SMLM to overcome the challenges posed by RI mismatches in deep tissue imaging. By evaluating various RI matched buffers, with a particular focus on the CUBIC-R+ based buffer, we demonstrated their efficacy through high-quality, dual-color 3D SMLM images across various depths in both cultured cells and sectioned tissue samples. The buffer’s ability to match the RI of oil and effectively clear tissue samples demonstrate the potential of RI matching as a straightforward and effective strategy to extend the imaging depth of SMLM, thereby enhancing its applicability in biological and biomedical research.

This research underscores the capability of RI matched buffers to facilitate high-quality, high-resolution imaging of biological structures at depths previously unachievable with standard SMLM approaches. For the first time, we have demonstrated an SMLM imaging buffer which is capable of tissue clearing and matched to the refractive index of immersion oil. We found that the duty cycle and survival fraction which is crucial for maintaining single molecule blinking images did not change much after adding the high RI liquid to the WI-based imaging buffer. Although we found the reduced emitted photon number of dyes when using CUBIC-R+ to match the RI of oil, the total collected number of photons within cellular environments in CUBIC-R+ based buffer is comparable with that in WI-based buffer due to higher photon collection efficiency in high RI buffer when a high NA objective was employed. The successful application of these buffers across multiple imaging scenarios paves the way for their broader adoption in biomedical research. In the future, the mechanism of the fluorophores within the high RI imaging buffer needs to be further elucidated. The optical properties of more dyes need to be evaluated if more than two color imaging is necessary. Ultimately, the findings advocate for the integration of RI-matching techniques into routine SMLM protocols, potentially transforming the landscape of biological imaging and opening new avenues for exploring cellular and molecular processes at the nanoscale.

## Materials and Methods

### Refractive index matched imaging buffer

The common water-based buffer for SMLM imaging consisted of 50 mM Tris-HCl (pH 8.0), 10 mM NaCl, 10% (w/v) glucose, 0.5 mg/ml glucose oxidase (GOx, G7141, Sigma), 40 μg/ml catalase (CAT, C100, Sigma) and 35 mM cysteamine (MEA, 30070, Sigma). To prepare RI matched imaging buffers for the silicone oil/ oil objective, high RI reagents containing 2,2’-thiodiethanol (RI = 1.5215, TDE, 166782, Sigma), glycerol (RI = 1.4746, Gly, AR, Damao), sucrose (RI = ∼1.465 for 80% (w/v) solution (*30*), Suc, S0389, Sigma), CUBIC-R+ (mixture of antipyrine (RI = 1.5697, AP, D1876, TCI) and nicotinamide (RI = 1.466, B3, N0078, TCI)) and 3-pyridinemethanol (RI = 1.545, 3-PM, A10381, Thermo Fisher Scientific) were used to mix with the water-based SMLM buffer, respectively. For buffers with RI matched to silicone oil (RI = 1.406), we mixed the diluted high RI stock (38% (v/v) TDE, 55% (v/v) of 85% (v/v) Gly and 60% (v/v) of 80% (w/v) Suc) and the water-based buffer in a volume ratio of 75:25. Similarly, buffers with RI matched to oil (RI = 1.518) were prepared as above excepted that CUBIC-R+ (or 3-PM) buffer included 5% (w/v) glucose, 100 mM MEA and 83% (v/v) CUBIC-R+ (or 82% (v/v) 3-PM). Here, CUBIC-R+ stock was prepared by mixing 50% (w/w) AP, 30% (w/w) B3 and 20% (w/w) MilliQ water. The RIs were measured by using Abbe refractometer (WAY-2WAJ, Lichen, China).

### Fluorescent beads for evaluating the depth-dependent PSFs

Before sample preparation, high-precision 25-mm-round coverslips (1.5H, CG15XH, Thorlabs) were cleaned by sequentially sonicating in 1 M potassium hydroxide (KOH), Milli-Q water and ethanol. We incubated Tetraspeck beads (diameter 100 nm, T7279, Invitrogen) on coverslips with 0.1 M MgCl_2_. To compare the PSFs in different depths, two coverslips with beads were mounted on the sample holder, separated by layers of water or RI matched media (without GOx/CAT/MEA), and sealed by picodent twinsil. The different depths were obtained by adjusting the volume of media solution or adding a brain slice with fixed thickness of 20 μm as a spacer. Here, we defined the depth of beads on the bottom coverslip as 0, and calibrated the depth of beads on the top coverslip by the relative z displacement between the bottom and top. To acquire the 3D PSFs, we introduced astigmatism with a cylindrical lens, used a 640 nm laser to excite the beads’ samples and recorded z stacks from -1 μm to 1 μm with step of 20 nm. In addition, all the PSFs, acquired from different media and depths, were averaged from 15-20 beads, and the lateral and axial FWHM were determined for each PSF.

### Measurements of absorption and fluorescence spectra

For the prepared oil-index buffers (CUBIC-R+ and 3-PM), the corresponding absorption and emission spectra were measured on a UV-Vis (UV-2600, Shimadzu) or fluorescence (F-4700, Hitachi) spectrophotometer, respectively. For the measurements of fluorescence spectra, the two buffers were excited at different wavelengths (405, 488, 561 and 642 nm), respectively.

### Single-molecule fluorescence measurements

Two red-absorbing dyes, AF647 and CF680, with superior blinking behavior in water-based SMLM imaging buffer were selected as benchmarks to quantify the influence of the RI matched buffers on the photoswitching properties of dyes. The self-conjugated or commercial dye-labeled secondary antibodies, sab-AF647 and sab-CF680 (20817, Biotium), with ratios of ∼ 1 dye per antibody were diluted to 1-10 nM in MilliQ water containing 5 μM bovine serum albumin (BSA). Before adsorbing the diluted dye solutions on coverslips for 30-60 min, we used the plasma cleaner (PDC-MG, Mingheng, China) to further clean the high-precision coverslips. Subsequently, samples were washed with MilliQ water for three times with 5 min each. For the conjugation of sab-AF647, secondary antibodies (115-007-003, Jackson Immunoresearch) and AF647 dyes (A37573, Invitrogen) were mixed with a mole ratio of 1:3 in 0.1 M NaHCO_3_ buffer, and incubated for 1 h in the dark. The labeled antibodies were then purified using a NAP-5 column (17085302, GE Healthcare) and determined by a UV-Vis spectrophotometer.

For the comparison of the characteristics of dyes in RI matched buffers and those in water-based buffer, we recorded a series of sequentially raw images (30-ms exposure time and 60,000 frames) by using a 640 nm laser (1 kW/cm^2^) to excite samples. As shown in Fig S14, we could directly separate the specific dye signal from the background by filtering out the dim signal. Thus, from the raw time-lapse image sequences, we could extract fluorescence time traces of individual dye molecules as follow. Briefly, we began with the detection and localization of fluorescent signals from individual dyes by SMAP software (*31*). The too wide or too dim localizations were rejected by setting the PSF width (< 160 nm), photon count (> 800) and x-y precision (0-10 nm). We then grouped localizations into individual clusters via hierarchical agglomerative clustering with a cutoff distance of 100 nm, and regarded the localization clusters with FWHM below 30 nm as signals from single dyes. Only clusters from single dyes were selected for the extraction of fluorescence time traces. Usually, four parameters, number of photons detected per switching event, number of switching cycles, on-off duty cycle (DC, fraction of time in the on state) and survival fraction (SF, fraction of dyes not photobleached), were measured to evaluate the photoswitching properties of dyes (*23*). Here, photons per switching event was determined by summing the photons in consecutive frames for each molecule. DC was measured within a sliding window of 100 s and SF was calculated within the initial 700 s. The equilibrium DC (or SF) were determined by the mean value between 400-600 s (or at 400 s).

### Cell culture and sample preparation

U2OS-Nup96-SNAP (300444, Cell Line Services) were grown in DMEM (10569, Gibco) containing 1×MEM NEAA (11140-050, Gibco), 10% (v/v) fetal bovine serum (FBS, 10099-141C, Gibco), 100 U/ml penicillin and 100 μg/ml streptomycin (PS, 15140-122, Gibco). COS-7 cells (100044, BNCC) were also grown in DMEM with FBS and PS. All cells were cultured in a humidified atmosphere with 5% CO_2_ at 37 °C and passaged every two or three days.

For super-resolution imaging of nuclear pore complex, U2OS-Nup96-SNAP cells were cultured on the clean coverslips for 2 d with a confluency of ∼80%. We labeled Nup96-SNAP with SNAP-tag ligand O^6^-benzylguanine conjugated AF647 (BG-AF647, S9136S, New England Biolabs) as described else (*26*). Briefly, cells were prefixed in 2.4% (w/v) paraformaldehyde (PFA) for 30 s, permeabilized in 0.4% (v/v) Triton X-100 (T8787, Sigma) for 3 min and further fixed in 2.4% PFA for 30 min. Then, cells were quenched in 0.1 M NH_4_Cl for 5 min and washed twice with PBS. To decrease unspecific binding, cells were blocked for 30 min with Image-iT FX Signal Enhancer (I36933, Invitrogen). Subsequently, cells were incubated in dye solution (1 μM BG-AF647, 1 mM DTT and 0.5% BSA in PBS) for 1 h, and washed 3 times in PBS for 5 min each to remove excess dyes. Lastly, cells were postfixed with 4% PFA for 10 min, washed with PBS 3 times and stored at 4 °C until imaged in imaging buffer.

For dual-color imaging of microtubules and mitochondria, COS-7 cells were labeled with primary antibodies (mouse anti-β-tubulin, 1:600, T4026, Sigma; rabbit anti-Tom20, 1:1000, ab78547, Abcam) and corresponding secondary antibodies conjugated AF647 (anti-rabbit, 1:2000, A21245, Invitrogen) or CF680 (anti-mouse, 1:250, 20817, Biotium) for 2 h at room temperature, respectively. Before labeling, cells were firstly fixed with 3% PFA and 0.1% (v/v) glutaraldehyde (GA) in cytoskeleton buffer (CB pH 6.1, 10 mM MES, 5 mM glucose, 150 mM NaCl, 5 mM MgCl_2_, and 5 mM EGTA) for 15 min. Then, cells were quenched in PBS buffer with 0.5% (w/v) NaBH_4_ for 7 min, permeabilized with 0.3% IGEPAL CA-630 (I8896, Sigma) and 0.05% Triton X-100 for 3 min, and blocked with 3% BSA for 1 h. Finally, cells were postfixed with 3% PFA and 0.1% GA for 10 min, and stored in PBS at 4 °C until imaged in imaging buffer.

Before imaging in CUBIC-R+ or 3-PM buffers, samples were successively immersed in the corresponding diluted solutions (10%, 25%, and 50% (v/v), respectively, in PBS, 5-10 min each) to exchange the water with CUBIC-R+ or 3-PM in a stepwise manner.

### Brain slice and MOG staining

All animal experiments were conducted in compliance with the protocols approved by the Animal Care Committee at Southern University of Science and Technology (SUSTech), China. To prepare brain sections, adult mice (8-10 weeks) were anaesthetized with isoflurane and transcardially perfused with 4% PFA. The brains were then submerged and postfixed in 4% PFA at 4 °C overnight, cryoprotected and dehydrated with 15% (w/v) sucrose for one day and 30% sucrose for a second day. Thereafter, the brain tissues were embedded in Tissue-Tek OCT (4583, Sakura, Japan) for 12 h, frozen with dry ice, and stored at −80 °C. Coronal or sagittal sections with a thickness of 20 or 30 μm were cut on a cryostat (Minux FS800, RWD).

For MOG staining, the coronal sections were first rinsed with PBS three times for 10 min each, and then blocked with 5% BSA and 0.3% Triton X-100 in PBS (blocking solution) for 1 h. Subsequently, slices were labeled by mouse anti-MOG (1:250, MAB5680, Sigma) in blocking solution overnight at 4 °C. The next day, slices were washed three times with PBS, and then incubated with secondary goat anti-mouse antibodies conjugated AF647 (1:500, A21235, Invitrogen) in blocking solution for 2 h. Finally, slices were washed three times with PBS and stored in PBS at 4 °C before imaging. To fix the brain slice on a precision coverslip, mixture of 0.5% gelatin and 0.05% chromium potassium sulfate was deposited on the tissue-coverslip layer, and dried at 60 °C for 5 min (*32*).

### Optical Setup

SMLM imaging were conducted on a custom-built microscope equipped with a silicone-oil or oil immersion objective (100x, NA 1.35, UPLSAPO100XS, Olympus; 100x, NA 1.5, UPLAPO100XOHR, Olympus), four lasers (405/488/561/640 nm), a main dichroic mirror (ZT405/488/561/640rpcxt-UF2, Chroma), a main emission filter (EF, NF03-405/488/561/635E-25, Semrock) and a sCMOS camera (ORCA-Flash4.0 V3, Hamamatsu). A band-pass EF (ET700/100m, Chroma) was inserted in the imaging optical path for further excluding residual laser light. The z focus was stabilized by a closed-loop system, which coupled the reflected signal of a 785 nm laser on the coverslip and the detected signal on a quadrant photodiode (SD197-23-21-041, Advanced Photonix) with the piezo objective z stage (P-726.1CD, Physik Instrumente).

For 3D SMLM imaging, we used a cylindrical lens to introduce astigmatism PSF, and acquired z stacks of TetraSpeck microspheres (T7279, Invitrogen) on a coverslip to generate the experimental PSF model (*33*). For ratiometric dual-color imaging of AF647 and CF680, we used a 680LP dichroic mirror (T680lpxxr, Chroma) to split the emitted fluorescence, which was then recorded on two parts of the sCMOS camera. The global multi-channel experimental PSF model was also generated based on the z stacks of beads on a coverslip (*34*). The color of single molecules was assigned by the ratio of the intensities in each channel. Here, samples were excited with a 640 nm laser (1-3 kW/cm^2^), and concurrently activated with a 405 nm laser (0-1 W/cm^2^). For Nup96 and MOG proteins, 80,000 frames with 20-ms exposure time were recorded to reconstruct the SMLM images. For β-tubulin and Tom20 proteins, 100,000 frames were recorded with exposure time of 20 ms.

## Data analysis

The raw single- or dual-color data were fitted or global fitted with SMAP software written in MATLAB as described previously (*26, 31, 34*). For raw data acquired from water-based SMLM imaging buffer (RI = 1.352), the fitted z localizations for cells or tissue slices were corrected with a RI mismatch factor (F) which was calculated as Eq.1 (*35*).

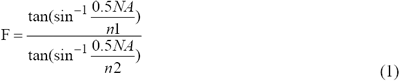

where NA is numerical aperture of the objective (1.35 for silicone-oil objective, and 1.5 for oil objective); n1 and n2 represent the RIs for the objective oil and the sample medium, respectively. Thus, the F values for silicone oil (RI = 1.406) and standard oil (RI = 1.518) objectives were measured as 0.95 and 0.85, respectively.

For sample drift, the x, y and z positions were corrected based on redundant cross-correlation algorithm. We regarded the localizations spanning consecutive frames (within a distance of 35 nm) as a single molecule, and thus merged into one localization. To reject dim localizations and bad fits, thresholds were set based on the type of data. 3D data of Nup96 were filtered by x-y precision (0-5 nm), z precision (0-30 nm), photon count (> 800), relative log-likelihood (> -2.5) and frames (1000-79000).

3D dual-color data of tubulin and Tom20 were filtered by x-y precision (0-10 nm), z precision (0-30 nm), photon count (> 800), relative log-likelihood (> -2.5) and frames (> 1000).

3D data of MOG were filtered by x-y precision (0-20 nm), z precision (0-45 nm), photon count (> 800), relative log-likelihood (> -2.5) and frames (> 1000).

To quantify the 3D data of Nup96, we automatically segmented NPCs in SMAP as depicted previously (*24*). Briefly, we convolved the super-resolution image with a Gaussian ring kernel (diameter 85-125 nm) and then extracted the candidate NPCs by rejecting the local maxima less than a threshold (0.06). To further clean up the aberrant structures, we fitted the localizations in each candidate to a circle and excluded the candidates with too small (< 40 nm) or too large (> 70 nm) radius. Next, we re-fitted the localizations to a circle with fixed radius (55 nm), and discarded the structures where more than 25% of localizations were within 40 nm of the center or more than 40% of localizations were further away than 70 nm. Moreover, we discarded the candidates with less than 10 localizations to ensure sufficient sampling.

All the segmented structures were then characterized by parameters including the distance between cytoplasmic and nucleoplasmic rings and effective labeling efficiency (ELE, fraction of the eight visible corners in each NPC).

## Author contributions

Conceptualization: Y. L. Experimental design: L. Z., Y. L. Data acquisition: L. Z., W. S. and M. L. Data analysis: L. Z., J. C., K. F. Simulation: S. F. Project supervision: Y. L. Writing: L. Z. and Y. L.

## Data and materials availability

The data that support the findings of this study are available from the corresponding author upon reasonable request.

