## Supplementary material for "High Refractive Index Imaging Buffer for Dual Color 3D SMLM Imaging of Thick Samples": Figs. S1 to S14

Lulu Zhou *et al.*

**This PDF file includes:**

Figs. S1 to S14

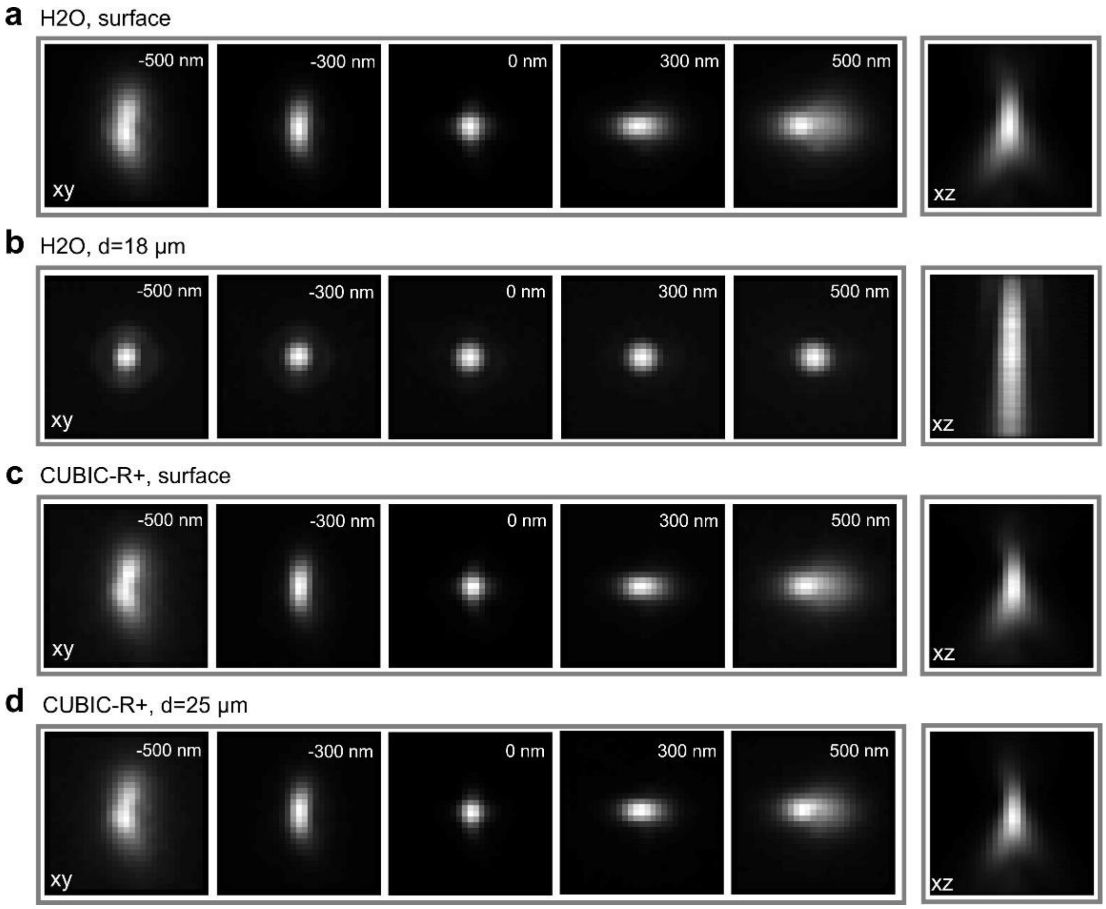


**Fig. S1. Comparison of the depth-dependent 3D PSFs in MilliQ water and RI matched medium (CUBIC-R+) with oil objective.** (a, b) Representative images of astigmatic PSFs in water at the bottom surface (a) and the depth of 18 µm (b). (c, d) Representative images of PSFs in CUBIC-R+ at the bottom surface (c) and the depth of 25 µm (d). 3D PSFs at bottom surface show no significant difference between the two media, whereas RI mismatch medium (H2O) introduced a significant spherical aberration for the deep-depth PSF.

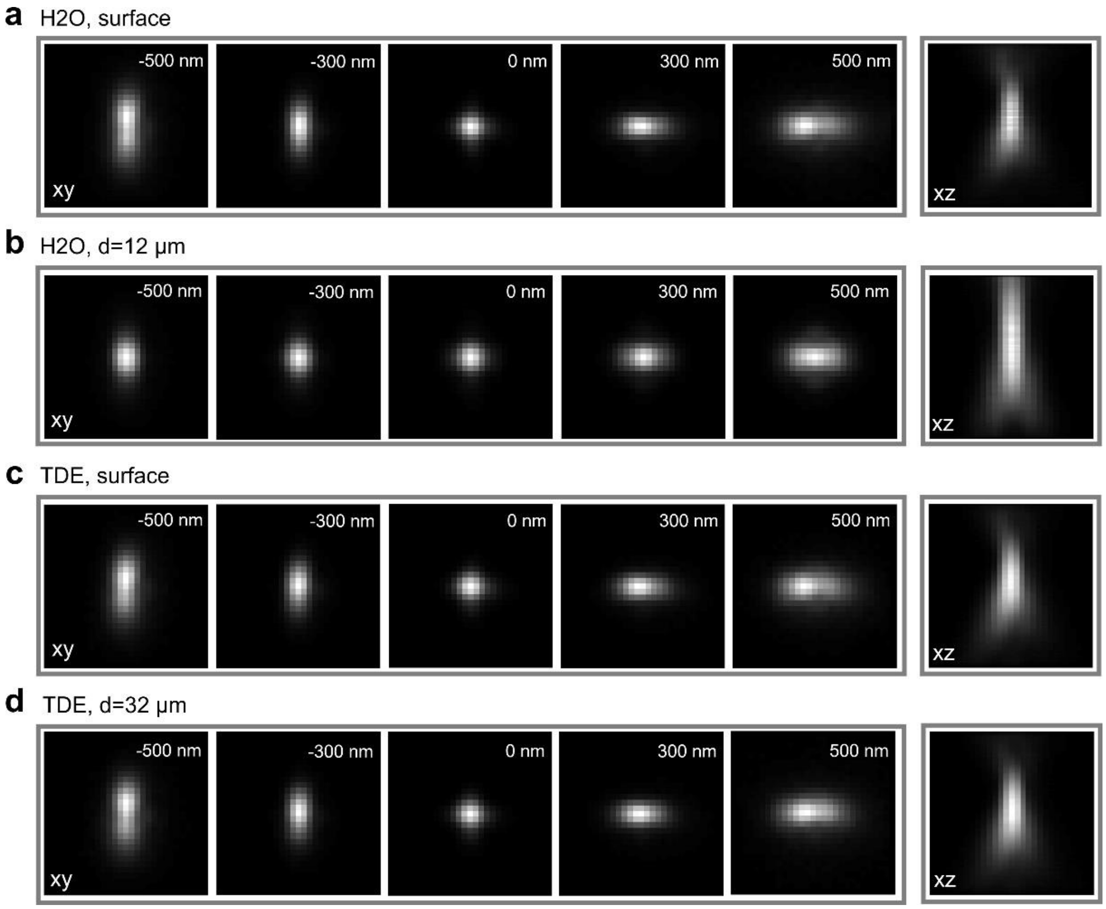


**Fig. S2. Comparison of the depth-dependent PSFs in MilliQ water and RI matched medium (TDE) with silicone oil objective.** (a, b) Representative images of 3D astigmatic PSFs in water at the bottom surface (a) and the depth of 12 µm (b). (c, d) Representative images of PSFs in TDE at the bottom surface (c) and the depth of 32 µm (d).

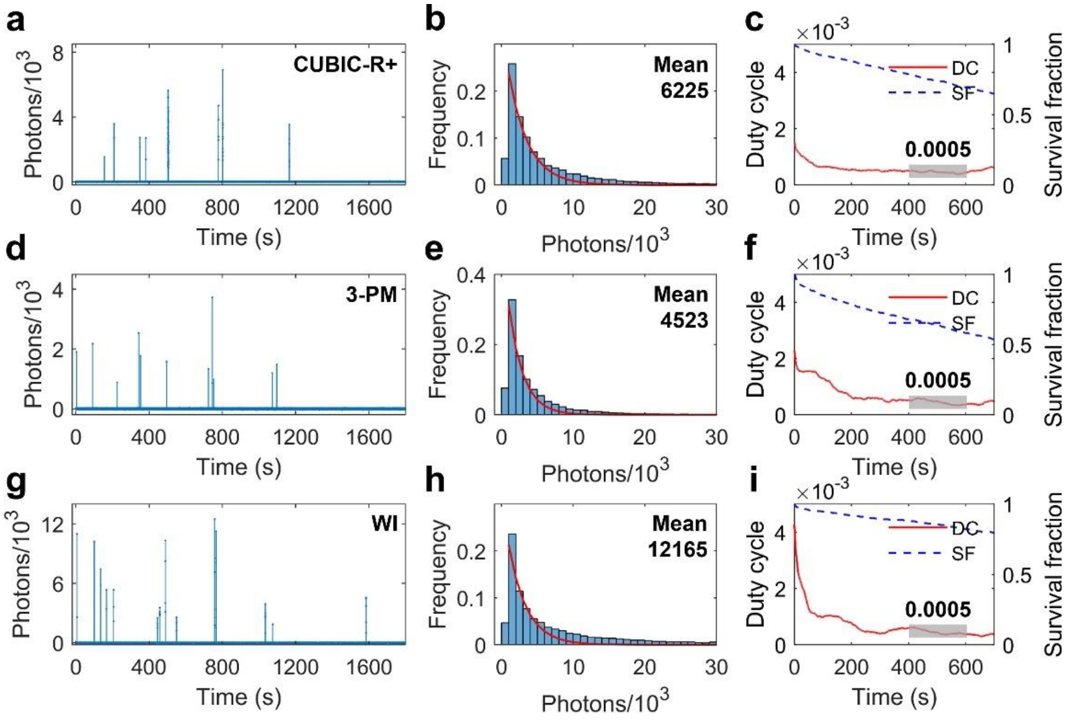


**Fig. S3. Comparison of the switching properties for AF647 in oil-index and water-index (WI) buffers.** (a-c) Representative single-molecule fluorescence time trace (a), histogram of photon number distribution (b) and duty cycle plot along with survival fraction as the function of time (c) for AF647 in CUBIC-R+ buffer. (d-f) Same as (a-c) but for AF647 in 3-PM buffer. (g-i) Same as (a-c) but for AF647 in WI buffer.

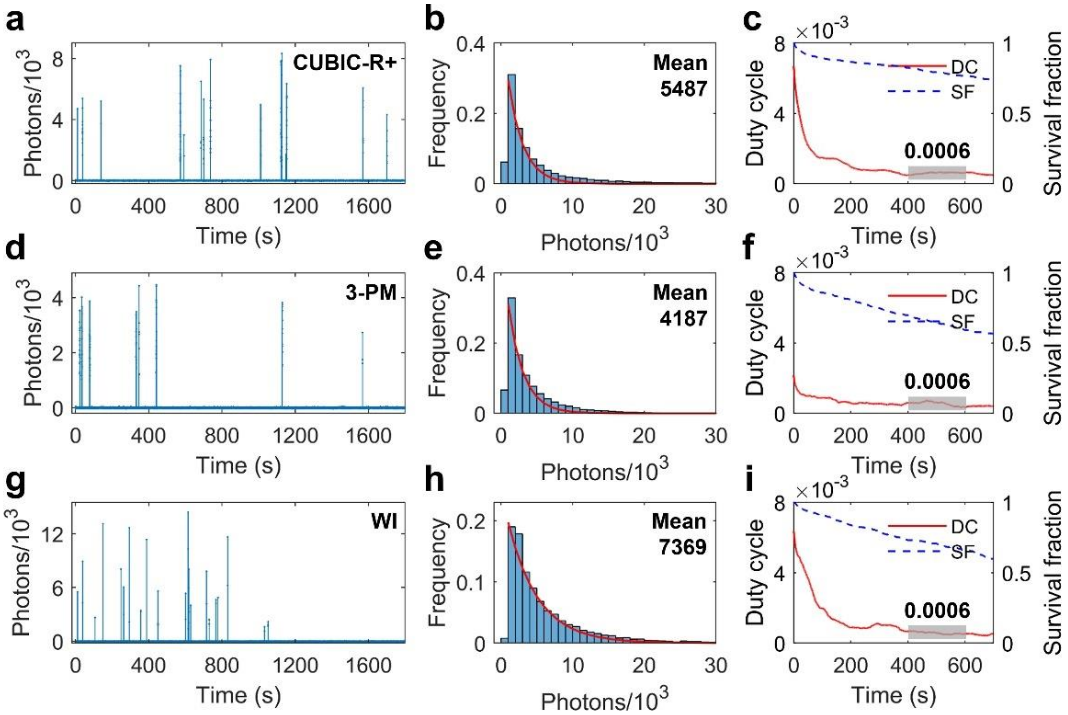


**Fig. S4. Switching properties of CF680 in oil-index and WI buffers.** (a-c) Representative single-molecule fluorescence time trace (a), histogram of photon number distribution (b) and duty cycle plot along with survival fraction as the function of time (c) for CF680 in CUBIC-R+ buffer. (d-f) Same as (a-c) but for CF680 in 3-PM buffer. (g-i) Same as (a-c) but for CF680 in WI buffer.

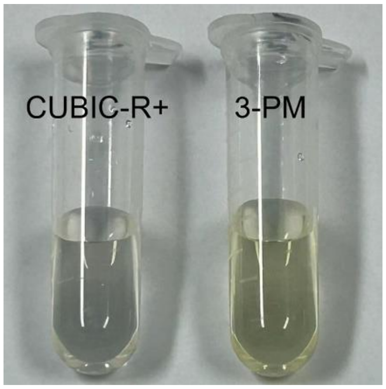


**Fig. S5. Comparison of the appearance of CUBIC-R+ and 3-PM buffers for SMLM imaging.**

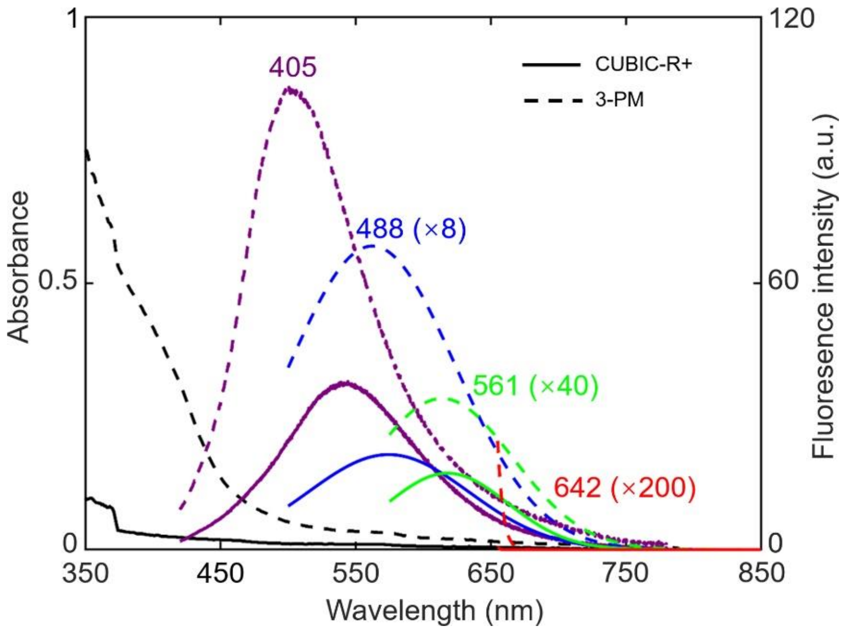


**Fig. S6. Comparison of the absorption and fluorescence spectra of CUBIC-R+ and 3-PM buffers for SMLM imaging.** The solid and dashed lines indicate the CUBIC-R+ and 3-PM buffers, respectively. The black lines show the absorption spectra. The violet, blue, green and red lines represent the emission spectra upon the excitation of 405, 488, 561 and 642 nm, respectively. For clarity, the blue, green and red fluorescence spectra were scaled by 8×, 40× and 200×, respectively.

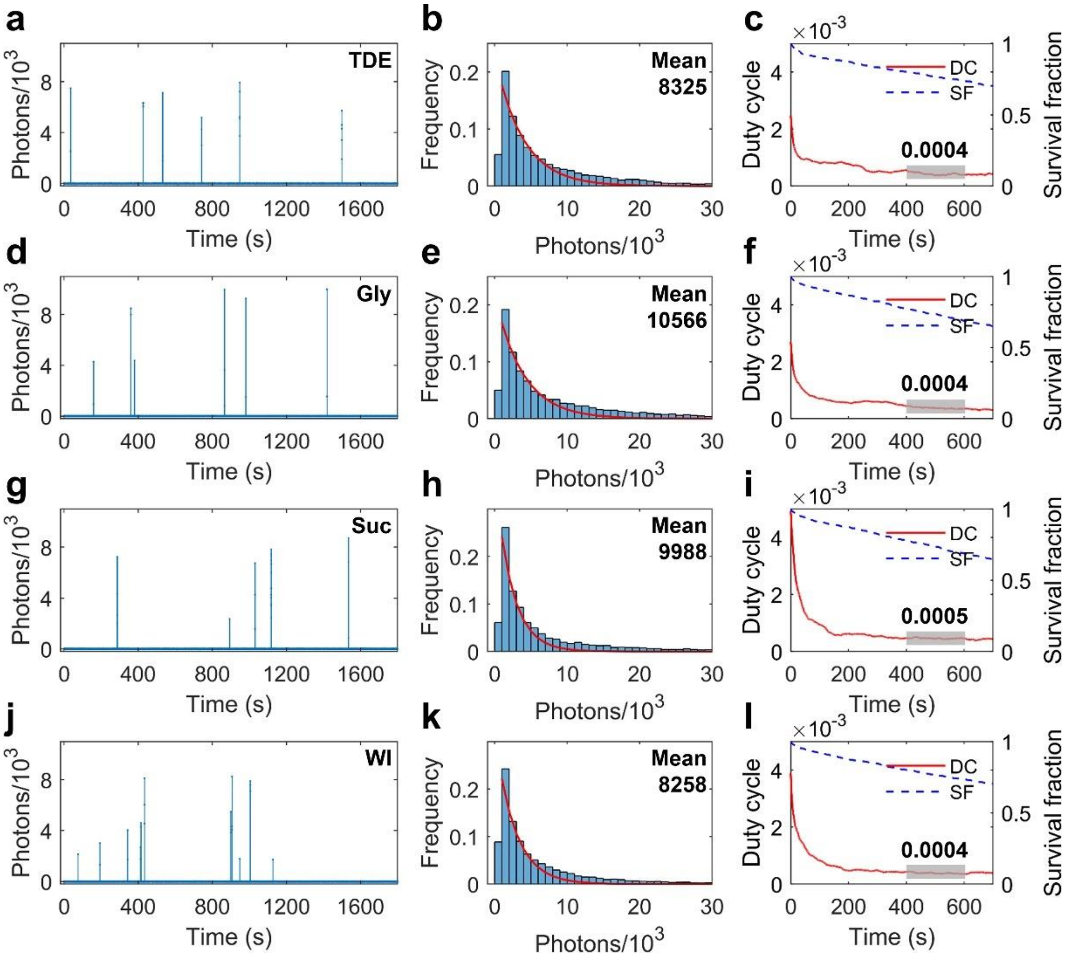


**Fig. S7. Switching properties of AF647 in water-based imaging buffer and silicone oil-index buffers.** (a-c) Representative single-molecule fluorescence time trace (a), histogram of photon number distribution (b) and duty cycle plot along with survival fraction as the function of time (c) for AF647 in TDE buffer. (d-f) Same as (a-c) but for AF647 in the glycerol buffer. (g-i) Same as (a-c) but for AF647 in the sucrose buffer. (j-l) Same as (a-c) but for AF647 in WI buffer.

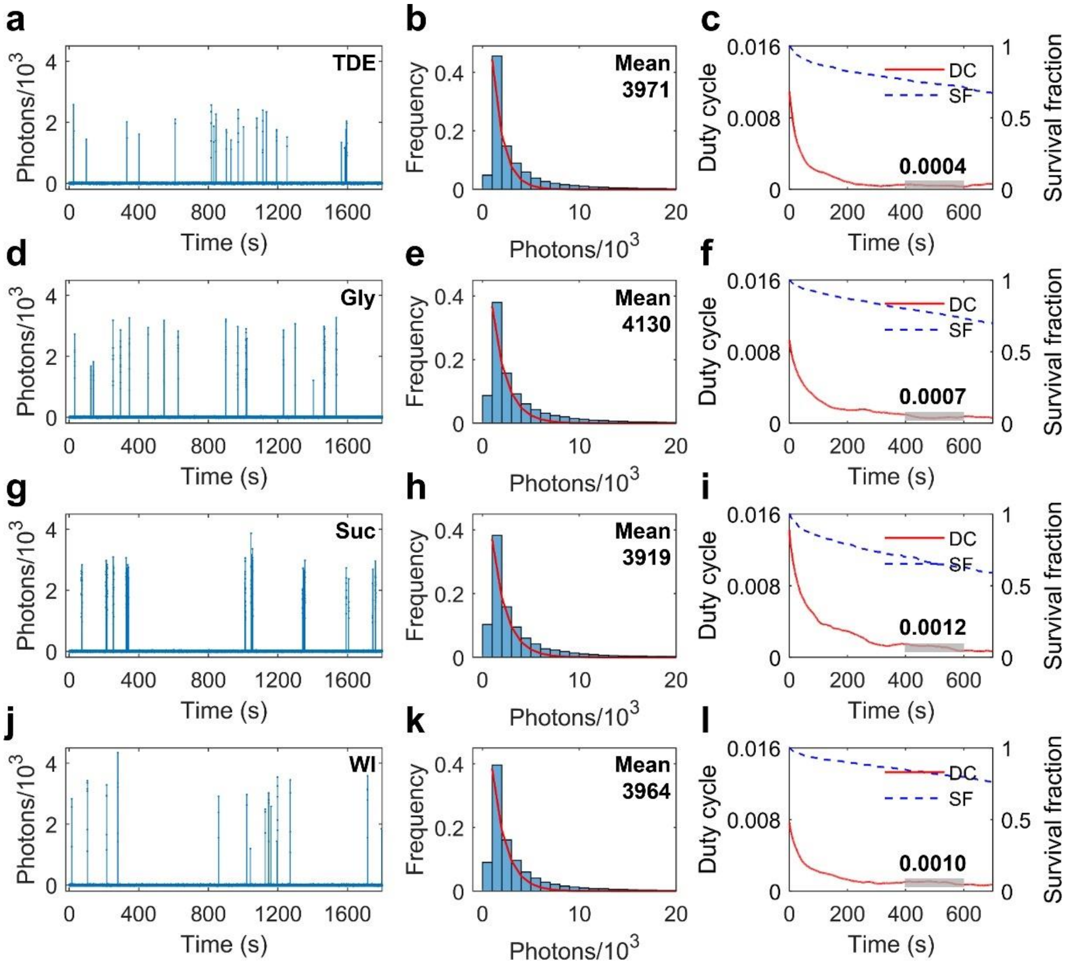


**Fig. S8. Switching properties of CF680 in water-based imaging buffer and silicone oil-index buffers.** (a-c) Representative single-molecule fluorescence time trace (a), histogram of photon number distribution (b) and duty cycle plot along with survival fraction as the function of time (c) for CF680 in TDE buffer. (d-f) Same as (a-c) but for CF680 in the glycerol buffer. (g-i) Same as (a-c) but for CF680 in the sucrose buffer. (j-l) Same as (a-c) but for CF680 in WI buffer.

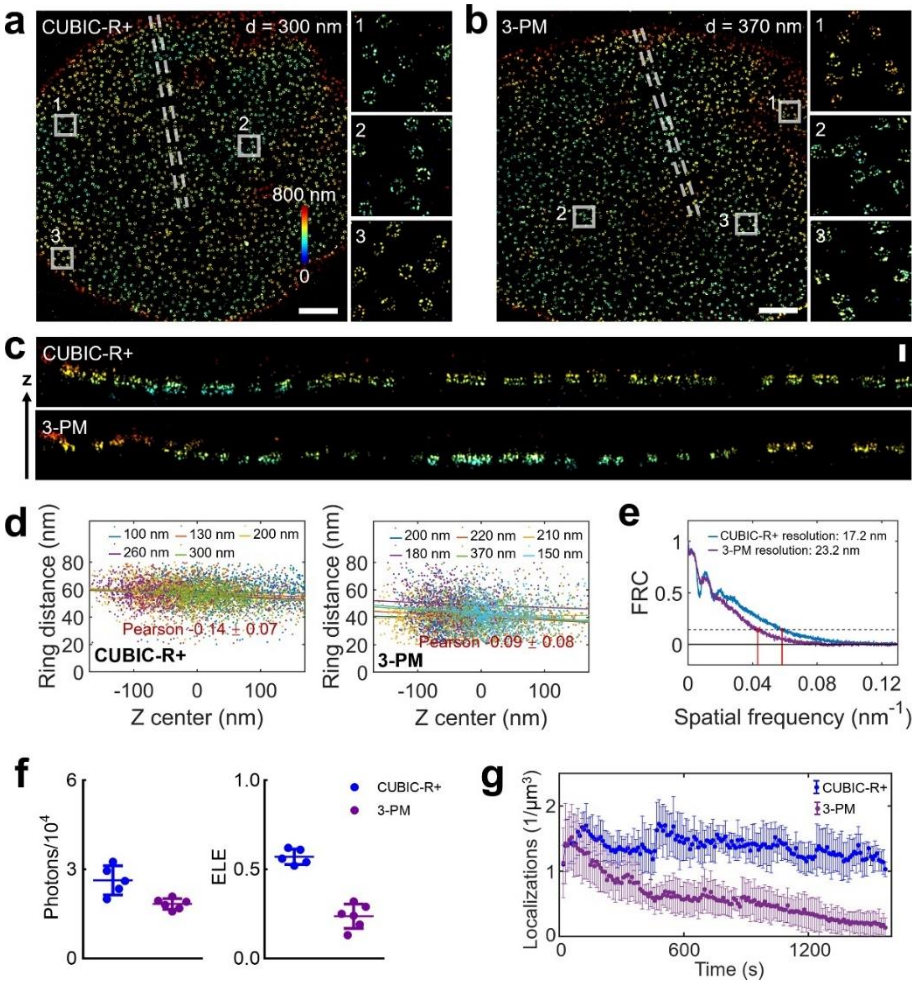


**Fig. S9. 3D SMLM imaging of Nup96 proteins on the lower nuclear envelope in oil-index buffers.** (a, b) Representative images of Nup96 labeled by AF647 in CUBIC-R+ (a) and 3-PM (b) buffers. The right panels show zoomed images for regions in white, and the color bar represents the z position. (c) The cross-sectional view images of the white dashed regions in (a, b). (d) Plots of distance between two rings versus the z position for Nup96 in different depths and buffers. (e) FRC curves of Nup96 in regions corresponding to the left panels of (a, b). (f) Scatter plots of the photon number (left) and ELE (right) for Nup96 in oil-index buffers. Each symbol corresponds to one cell nucleus. Horizontal bars and errors show the means and SD, respectively. (g) Comparing the number of localizations per μm$^3$ observed in CUBIC-R+ with 3-PM. Error bars show the SD. Scale bars: 2 μm (a, b), 200 nm (c).

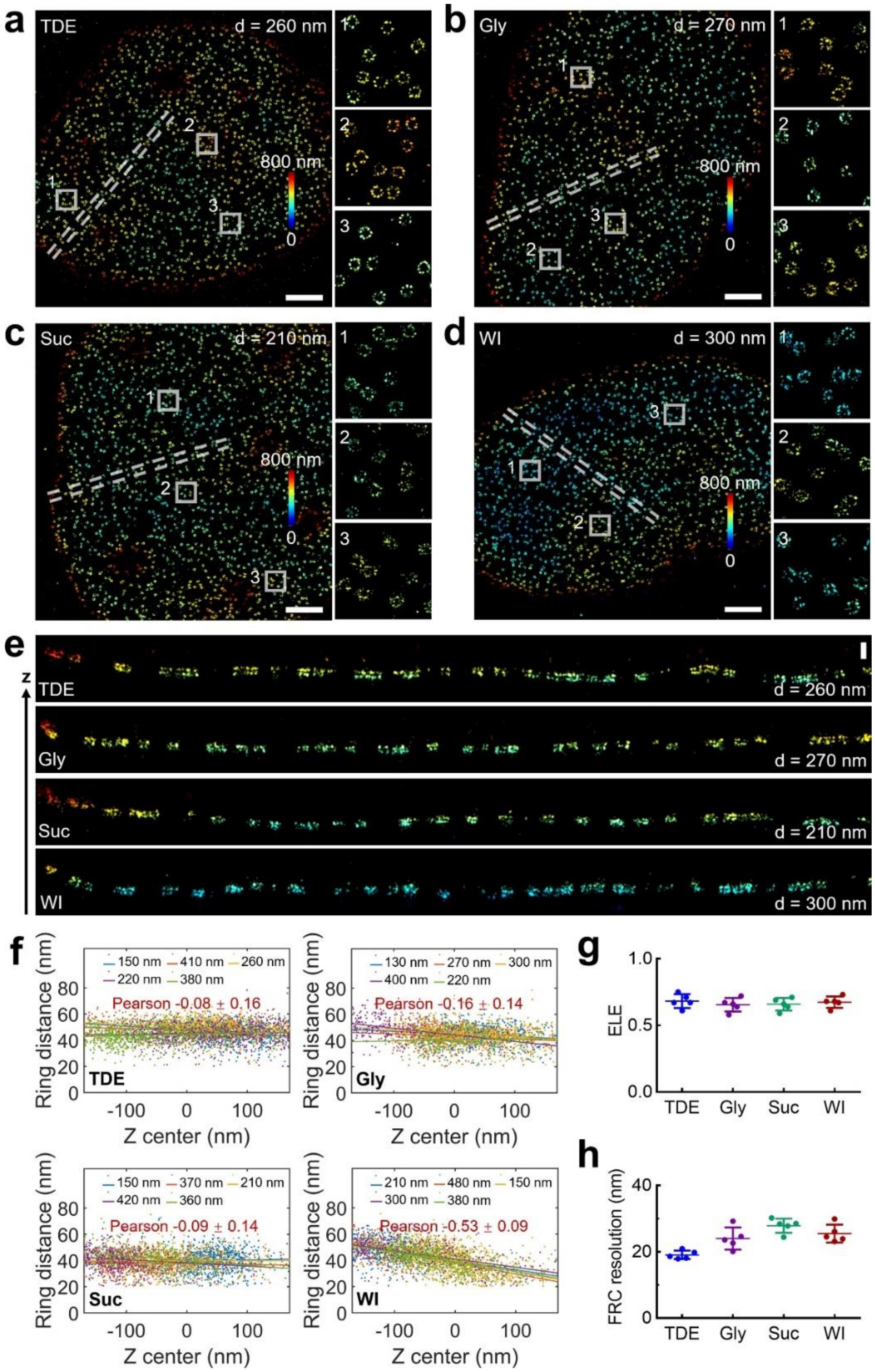


**Fig. S10. 3D SMLM imaging of Nup96 proteins on the lower nuclear envelope in silicone oil-index buffers.** (a-d) Representative images of Nup96 labeled by AF647 in TDE (a), glycerol (b), sucrose (c) and WI (d) buffers. The right panels show zoomed images for regions in white, and the color bar represents the z position. (e) The cross-sectional view images of the white dashed regions in (a-d). (f) Plots of distance between two rings versus the z position for Nup96 in different depths and buffers. (g, h) Scatter plots of the ELE (g) or FRC resolution (h) for Nup96 in silicone oil-index buffers. Each symbol corresponds to one cell nucleus. Horizontal bars and errors show the means and SD, respectively. Scale bars: 2 μm (a-d), 200 nm (e).

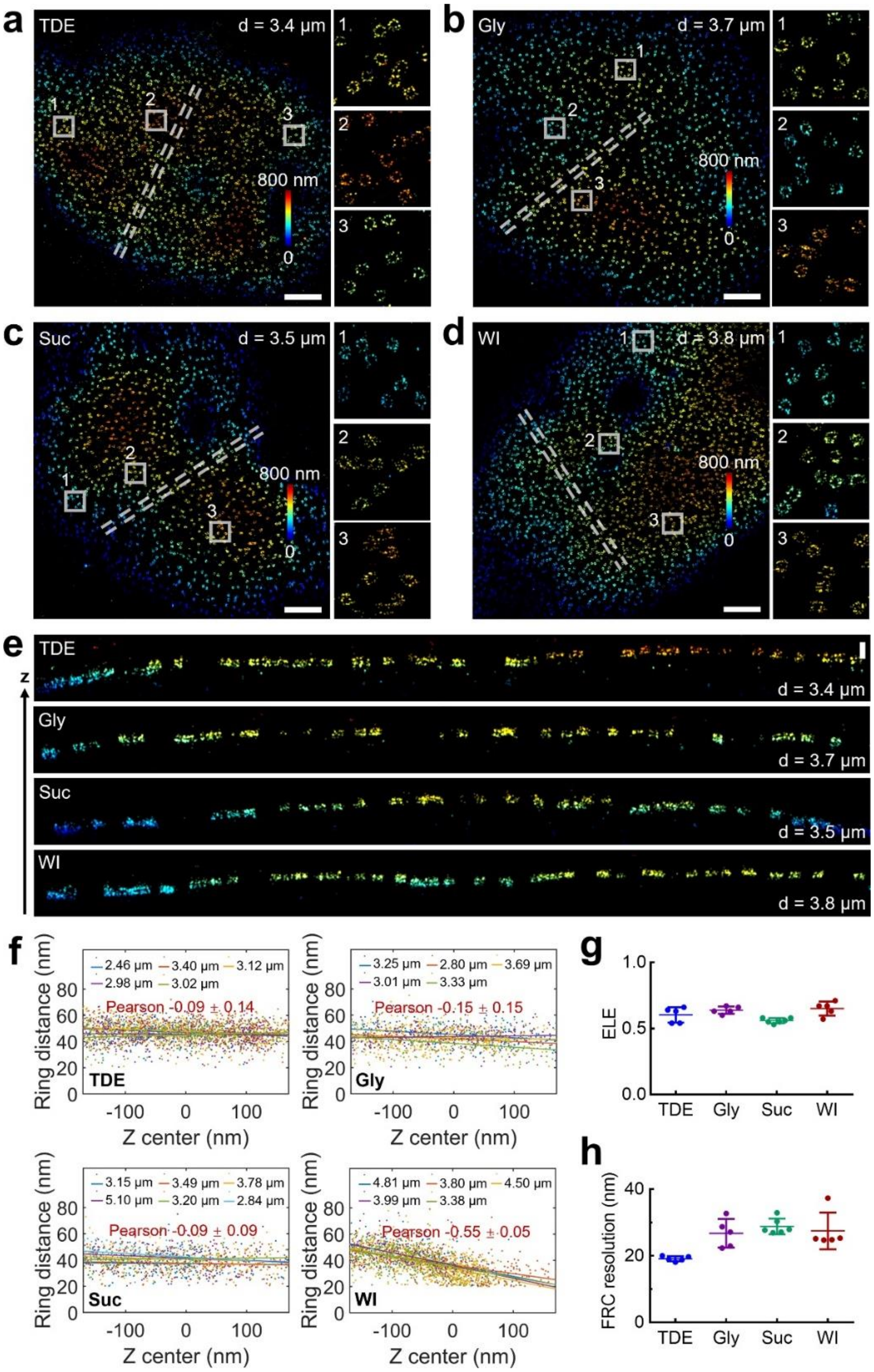


**Fig. S11. 3D SMLM imaging of Nup96 proteins on the upper nuclear envelope in silicone oil-index buffers.** (a-d) Representative images of Nup96 labeled by AF647 in TDE (a), glycerol (b), sucrose (c) and WI (d) buffers. The right panels show zoomed images for regions in white, and the color bar represents the z position. (e) The cross-sectional view images of the white dashed regions in (a-d). (f) Plots of distance between two rings versus the z position for Nup96 in different depths and buffers. (g, h) Scatter plots of the ELE (g) or FRC resolution (h) for Nup96 in silicone oil-index buffers. Each symbol corresponds to one cell nucleus. Horizontal bars and errors show the means and SD, respectively. Scale bars: 2 μm (a-d), 200 nm (e).

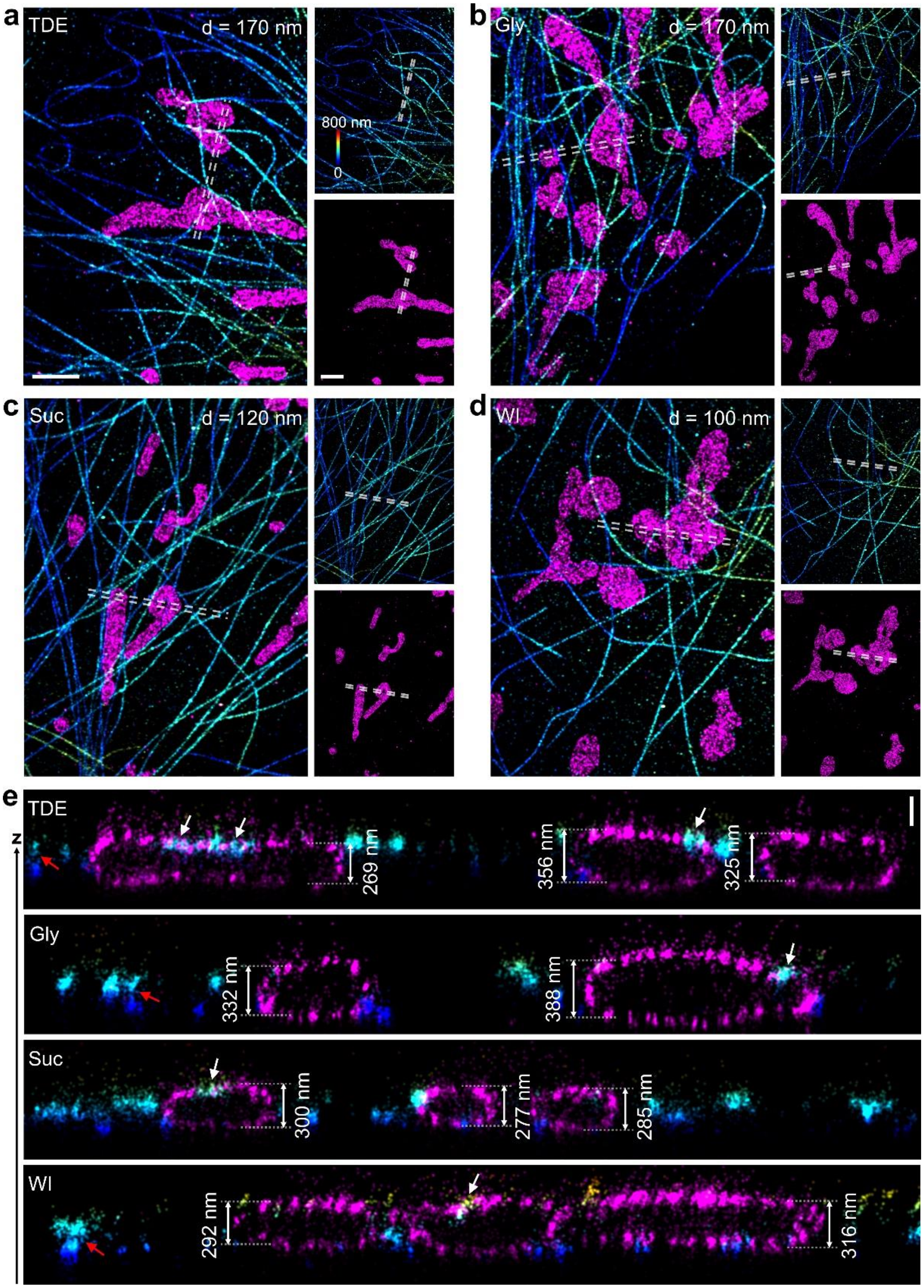


**Fig. S12. Dual-color 3D SMLM imaging of microtubules and mitochondria slightly above the basal membrane of COS-7 cells in silicone oil-index or WI buffers.** (a-d) Representative images of microtubules (anti-β-tubulin/CF680) and the outer mitochondrial membrane (anti-Tom20/AF647) in TDE (a), glycerol (b), sucrose (c) and WI (d) buffers. Color bar denotes the z position. (e) The side view of the white dashed regions in (a-d). The white arrows indicate the mitochondria-microtubule contacts, while the red arrows denote the different layers of microtubule filaments. Scale bars: 2 μm (a-d), 200 nm (e).

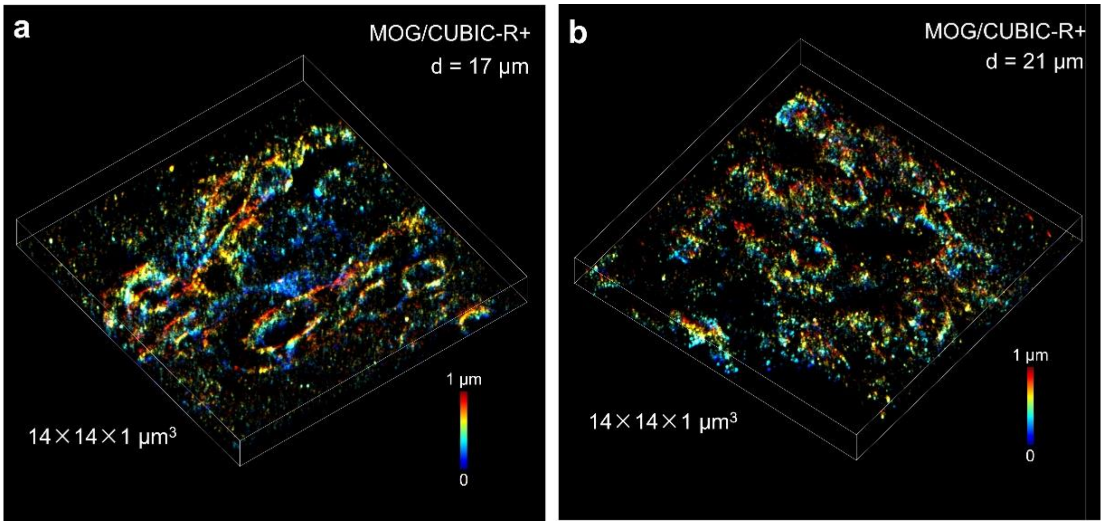


**Fig. S13. Deep 3D SMLM imaging of MOG proteins in mouse brain tissue slices with thickness of 30 µm.** (a, b) Representative 3D images of MOG labeled by AF647 in CUBIC-R+ buffer at different depths of 17 µm (a) and 21 µm (b). Color bars denote the z position.

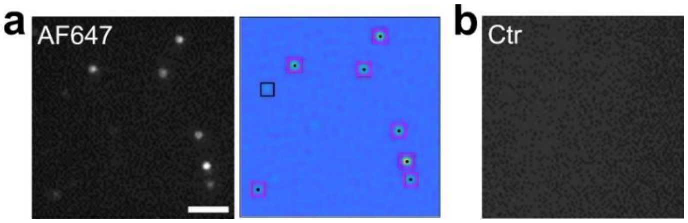


**Fig. S14. Distinguishing the single molecular signal from the background.** (a) Representative images showing individual molecules of AF647 on the coverslip under the WI imaging buffer. The black box in right image represents the dim signal which was discarded, whereas the magenta boxes represent the fitted localizations with enough high photons. (b) Representative image of the control group (without dyes on coverslip) showing no fluorescent signal.
